# Role of somatic HCN in epileptiform activity in subicular neurons

**DOI:** 10.1101/2024.07.26.604395

**Authors:** Monica Alfred, Sujit K. Sikdar

## Abstract

The subiculum, owing to its bursting nature and recurrent connections, plays a critical role in Temporal Lobe Epilepsy (TLE). Studying neuronal subtypes in the subiculum can elucidate the mechanisms underlying the patterning of epileptiform firing. We observed that epileptogenic 4AP-0Mg induced different patterns of epileptiform discharges in burst firing neurons and interneurons. Hyperpolarization-activated cyclic nucleotide-gated (HCN) channels regulate the intrinsic excitability of the neurons by governing the neuronal firing properties and membrane potential. To study the role of I_h_ (HCN currents) in epileptiform activity in subicular neurons, we modeled subicular HCN currents in the dynamic clamp that mimicked downregulation and overexpression observed in epilepsy-associated pathophysiology. Our results indicated that the burst firing neurons contribute to the epileptic firing characteristics due to HCN in the subiculum. We subsequently investigated the homeostatic modulation of HCN during the epileptiform activity in subicular burster cells. Our study is the first report showing I_h_ in rat subicular neurons during 4AP 0Mg-induced epileptogenic activity undergoes modulation on a time scale of a few minutes. Additionally, we observed that the changes in sag and chirp responses were persistent after the wash-out of 4AP-0Mg; thus, the changes appear irreversible. Our studies further showed that the neuronal excitability changes paralleled the changes in the HCN conductance during epileptogenesis. We conclude that a very rapid decline in somatic HCN function during epileptiform activity represents a previously unidentified mechanism of homeostatic dysfunction over a very short period, impeding the neuron’s ability to reestablish its regulatory processes in the subicular burster cells.

## INTRODUCTION

Temporal Lobe Epilepsy affects more than half of the epileptic population (1–2). The subiculum, a primary output gateway for the hippocampus, comprises burster neurons and exhibits less cell death in epileptic episodes, rendering it a promising region for investigating epileptogenesis. Subicular focal activity in TLE shows perturbations in the network excitation-inhibition (E-I) balance in which the burst firing population of neurons leads the initial dominance of excitation followed by the activity of interneurons that contributes to the delayed occurrence of inhibition (3). Not much has been studied on the mechanisms governing neuronal excitability and its transition to epileptic firing in these neuronal subtypes in the subiculum. Several studies over the last few years have shown increasing evidence for the potential role of HCN channels in the pathogenesis of epilepsy. Hyperpolarization-activated cyclic nucleotide-gated (HCN) channels are known to determine neuronal membrane potential and firing characteristics. Many results were derived from animal models (4–5), but further studies show dysfunctional HCN genetic mutations that are associated with epilepsy in human patients (6–8). We hypothesize that modulation of I_h_ in subicular neurons can regulate epileptogenic activity. This investigation can shed light on the mechanisms contributing to the different patterning of these subicular neurons towards epileptogenesis at the cellular level. The intriguing phenomenon of HCN in TLE has been observed in its tendency to upregulate and downregulate, depending on the physiological context (9). The dynamic clamp gives us a unique platform to explore these two physiological contexts by adding and subtracting HCN conductances in real time in the same neuron. In addition to these advantages, a dynamic clamp overcomes the limitations of selectivity issues of blocking HCN conductance pharmacologically as these blockers are also known to affect other voltage-gated ion channels like T-type Ca^2+^ (10) & L-type Ca^2+^ channels (11), and Na^+^ channels (12). Also, because of the lack of availability of I_h_ enhancers, it has been challenging to study the potential role of I_h_ upregulation in epileptogenesis, which can also be resolved with a dynamic clamp.

Our study reveals a differential response of I_h_ on epileptiform activity in burst firing and interneurons, with burst firing predominately influencing the epileptogenic firing patterns due to HCN in the subiculum. We suggest that the modulation of burst firing neurons by HCN during epileptiform activity could potentially regulate both the onset and the propagation of epileptogenesis. This modulation is crucial, as bursting plays a significant role in transmitting epileptiform discharges in TLE.

Studies are yet to be conducted observing the progression of HCN function during epileptogenic activity. In several reports, I_h_ has been known to play a primary role in both cellular homeostasis and epileptic mechanisms, suggesting a complex relationship between the two that needs further investigation (13–16). Here, we address the homeostatic modulation of HCN activity from the initiation of epileptogenic activity with its temporal progression, which has not been reported earlier. We observe a progressive decline in the HCN activity that occurs within a few minutes from the origin of the epileptiform activity. This is not only the first report of the homeostatic modulation during epileptiform activity occurring within minutes but also an initial report for demonstrating decreased HCN activity within a span of a few minutes in a neuron.

Several lines of evidence showed that seizures led to increased I_h_ in the hippocampal pyramidal neurons, including a depolarized shift of the activation curve and increased dendritic currents (9, 17, 18). Later studies showed that repeated epileptic episodes led to decreased I_h_ in several regions of the hippocampus and cortex (16, 19, 20). In this dual paradigm, our results corroborate with the observations from tissues from TLE patients that showed decreased I_h_ (21). Additionally, the acquired alterations in the HCN activity observed in our study were persistent post 4AP-0Mg washout. As alterations in HCN are associated with both decreased excitability and hyperexcitability (22), we sought to elucidate how epileptiform activity-induced irreversible changes in HCN function that our work demonstrates affect the overall excitability of the neurons.

## MATERIALS AND METHODS

All the experiments conducted in our study followed the guidelines provided by the Animal Ethics and Welfare Committee, Indian Institute of Science, Bangalore, India.

### Slice Preparation

Transverse hippocampal slices (350 µm) were obtained from male Sprague Dawley Rats of 18-24 days old. Brain dissection was done in oxygenated sucrose-based artificial cerebrospinal fluid (mM): 230 sucrose, 2.5 KCl, 1.25 NaH_2_PO_4,_ 25 NaHCO_3,_ 10 glucose, 3 sodium pyruvate, 0.5 CaCl_2_.2H_2_O, 7 MgCl_2_.6H_2_O using the Leica VT1000S vibratome. Hippocampal slices were transferred to incubation solution (mM): 125 NaCl, 25 glucose, 3 KCl, 1.25 NaH_2_PO_4_, 2 MgCl_2_.6H_2_O, 25 NaHCO_3_, 2 CaCl_2_.2H_2_O and 2 sodium pyruvate in distilled water (equilibrated with 95% O_2_ and 5% CO_2_, pH 7.4 with 300-310 mOsm) in an incubation chamber maintain at 34°C using a water bath.

### Electrophysiology

Patch pipettes (3-5 MΩ) were pulled from borosilicate glass capillaries (Harvard Apparatus) and filled with an internal solution (mM): 120 K-gluconate, 20 KCl, 10 HEPES, 5 Mg-ATP, 0.2 EGTA, 4 NaCl, 0.5 Na-GTP, 10 Na-phosphocreatinine (pH 7.3 and osmolarity of 290-300 mOsm). The hippocampal slices were perfused continuously with carbogen-saturated ACSF or normal ACSF (nACSF) (mM): 125 NaCl, 1.25 NaH_2_PO_4_, 25 NaHCO_3_, 3 KCl, 25 glucose, 1 MgCl_2_.6H_2_O, 2 CaCl_2_.2H_2_O (equilibrated with 95% O_2_ and 5% CO_2_, pH 7.4 with osmolarity of 290-300 mOsm). All the control experiments were done in nACSF. Neurons were visualized by a Dodt contrast video microscopy (BX51WI, Olympus) using (40x, 0.8 NA) objective lens. Somatic whole cell recordings were conducted in the fast current-clamp mode using the DAGAN BVC 700A amplifier (Dagan Corporation, MN, USA), Digidata 1440A Digitizer (Molecular Devices) filtered at 5kHz that were stored on disk at >20 kHz sampling frequency. Acquisition of data was done using WinWCP software (Strathclyde Electrophysiology Software, UK). Series resistance was compensated. Data from neurons exhibiting depolarized membrane potential than –55 mV were discarded.

Epileptiform activity was induced in vitro by perfusing the slices with 100µM 4-AP (4-Aminopyridine) in magnesium-free ACSF (4AP-0Mg) containing (in mM) 125 NaCl, 1.25 NaH_2_PO_4_, 25 NaHCO_3_, 3 KCl, 25 glucose, 2 CaCl_2_.2H_2_O (equilibrated with 95% O_2_ and 5% CO_2_, pH 7.4 with osmolarity of 200-310 mOsm). We used 100 μM 4AP-0Mg in the external solution to induce epileptiform activity in the hippocampal slices. 0Mg makes the epileptic model robust and last longer (23–24).

### RTXI

In our study, to examine the functional outcome of alterations in I_h_ in subicular neurons, we artificially manipulated HCN currents using the dynamic clamp in our current-clamp experiments. Our dynamic clamp consists of an amplifier (Axopatch 200B), data acquisition (NI-PCI 6221) and a computer using a real-time operating system that controls the entire set-up. The HCN conductance (G_h_) was simulated from the model used by van Welie, 2006 (and Magee, 1998) using the following equations:

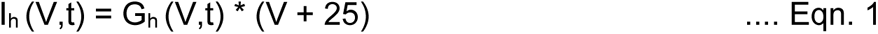

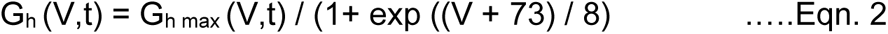

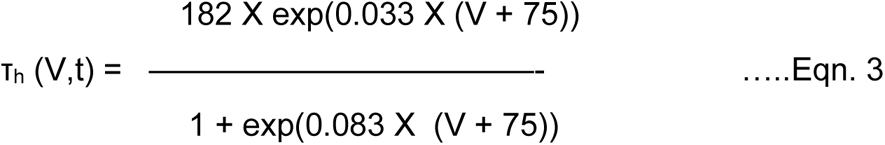

where I_h_: voltage and time dependent HCN current; G_h_: conductance of HCN channel; G_hmax_: maximal conductance of HCN channel; τ_h_: steady state kinetics of I_h_; V: membrane potential

### Analyses and Statistics

The analyses of the electrophysiological data were done in Igor Pro6 (Wave Matrix), MATLAB (Mathworks), GraphPad PRISM (GraphPad Software Inc., California, USA), and Clampfit 10.4 (Molecular Devices Inc., USA). The instantaneous spike frequency was estimated by calculating the inverse of the interspike interval for each time (t).

The sag ratio was calculated as the ratio of steady-state voltage deflection to peak voltage deflection.

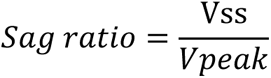

where, Vss=steady state voltage response of the hyperpolarizing current step pulse Vpeak=peak voltage response of the hyperpolarizing current step pulse

A chirp current stimulus of 0.1 nA amplitude and frequency spanning 0-20 Hz for 20 sec was used for analyzing subthreshold resonance. The impedance amplitude profile (ZAP) was estimated from the magnitude of the ratio of the fast Fourier transform (FFT) of the voltage response to the fast Fourier transform of the current stimulus. The resonance frequency F_R_ is the frequency that shows the peak impedance amplitude.

Shapiro-Wilks test was used to confirm if the data had a normal distribution. When the data did not fit into a normal distribution, the Wilcoxon matched-pairs signed rank test was used subsequently. ANOVA (two-way or three-way) test was used to see the statistical significance among and between different groups. The statistical significance was determined using student’s t-test for paired and unpaired groups. All data are represented as mean ± SEM, and n represents the number of samples. Statistical analyses were carried out in softwares like GraphPad Prism and MATLAB. P values < 0.05 were considered to be statistically significant.

## RESULTS

### 4AP 0Mg-Induced Epileptiform Firing is Higher in Interneurons than in Burst Firing Neurons

Burst firing neurons and fast-spiking interneurons (FSIN) were identified based on their electrophysiological properties. The firing characteristics of neurons were studied by applying a 1s long and 300 pA amplitude of depolarizing current step. Burst firing neurons exhibit a cluster of high frequency two or more action potentials that are followed by single action potential spikes (Fig. 1*Aa*). Interneurons elicit high frequency action potential with the firing rate >50Hz (Fig. 1*Ab*). The resting membrane potential (RMP) of the fast-spiking interneurons (FSIN) (–70.24 ± 3.16 mV, n=8) was more hyperpolarized in comparison to that of the burst firing neurons (–66.87 ± 1.19 mV, n=15).

**Figure 1.**
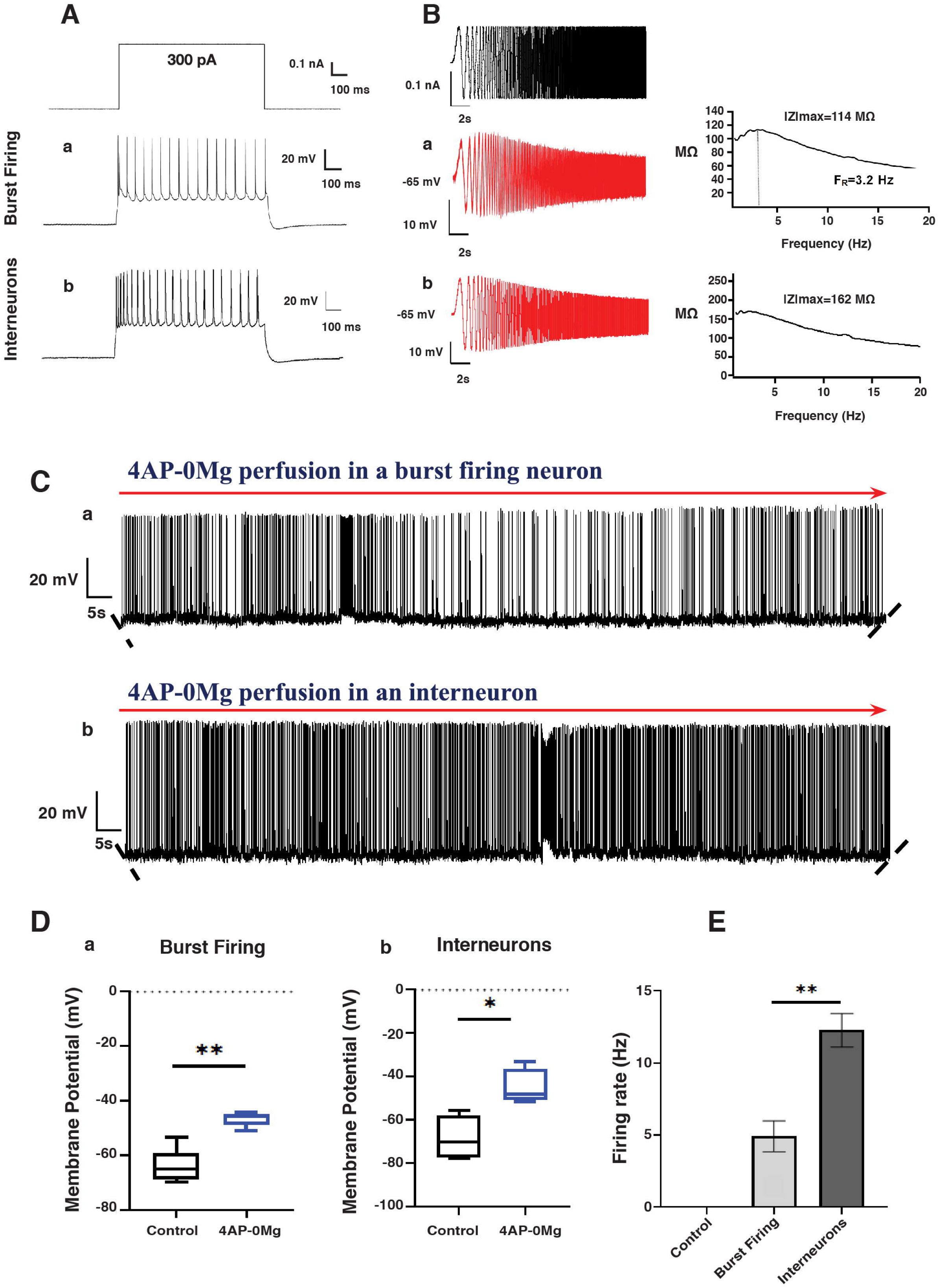
4AP-0Mg induced increased epileptiform firing in interneurons compared to burst firing neurons. (A) The neuronal response to 300 pA for 1 s *at –65 mV* in (a) burst firing neurons showed an initial burst (doublet) at the onset of the current step pulse that was followed by single action potentials (b) interneurons exhibited a high frequency action potential spike. (B) The voltage response to a sinusoidal stimulus of 0.1 nA at –65 mV of (a) a burst firing neuron and its impedance plot (right) depicting a theta resonance (b) an interneuron and the impedance plot (right) showing a low pass filtering. (C) A trace of voltage response for 3 mins with the bath application of 4AP-0Mg from (a) a burst firing neuron and (b) an interneuron. (D) Average plot displaying a shift in membrane potential after 4AP-0Mg induction in (a) burst firing neurons (control –64.14 ± 2.89 mV vs. 4AP-0Mg –46.49 ± 1.16 mV, n=5, **p<0.01, paired t-test) and (b) interneurons (control –67.96 ± 4.44 mV vs. 4AP 0Mg –44.41 ± 3.48 mV, n=5, *p<0.05, paired t-test). (E) Mean firing rate of epileptiform activity induced by 4AP-0Mg in burst firing neurons and interneurons as compared to control conditions (burst firing 4.92 ± 1.06 Hz vs. FSIN 12.27 ± 1.17 Hz, n=5, ** p=<0.01, unpaired t-test). Data represent mean ± SEM in all plots.

The resonance properties of these two classes of neurons were estimated using chirp current stimulus (25) of a constant amplitude of 0.1nA with increasing frequency (1Hz/second for 20s) applied at RMP –65 mV. The peak voltage response called as the resonance frequency, F_R_ was observed at theta frequency, 5.20 ± 1.0 Hz (n=5) (Fig. 1*Ba*) for burster cells, and their mean resonance amplitude was 167.53 ± 32.53 MΩ (n=5). Fast spiking interneurons showed a low pass voltage response to the chirp stimulus (Fig. 1*Bb*) 0.79 ± 0.11 Hz (n=7) and the mean resonance amplitude of 293.15 ± 21.68 MΩ (n=7).

We used 100 μM 4AP-0Mg in an external solution that induced epileptiform activity in the hippocampal slices within 5-10 minutes (23–24). Only those burst firing neurons and interneurons that showed the spontaneous occurrence of epileptiform activity (after the perfusion of 4AP-0Mg) were considered for further analysis. Analysis of the resting membrane potentials showed a significant depolarizing shift in membrane potential for burst firing neurons (control –64.14 ± 2.89 mV vs. 4AP-0Mg –46.49 ± 1.16 mV, n=5, **p<0.01, paired t-test) (Fig. 1*Da*) as well as for FSIN (control –67.96 ± 4.44 mV vs. 4AP 0Mg –44.41 ± 3.48 mV, n=5, *p<0.05, paired t-test) (Fig. 1*Db*) following the induction of epileptiform activity with 4AP-0Mg. We observed a significant difference in membrane potential in these two neuronal subtypes, which confirmed the establishment of 4AP-0Mg induced epileptiform activity. Under control conditions, the neurons were quiescent at RMPs –64.14 ± 2.89 mV for burst firing (n=5) and at RMPs –67.96 ± 4.44 mV for interneurons (n=5) with no spontaneous action potentials. However, bath application of 4AP-0Mg evoked epileptiform activity in both the types of neurons and showed a significant difference in the firing rate during epileptiform activity induced by 4AP-0Mg perfusion in burst firing neurons 4.92 ± 1.06 Hz and interneurons 12.27 ± 1.17 Hz (n=5 for each group, burst firing vs. FSIN, ** p=<0.01, unpaired t-test, Fig. 1*C a, b; E*).

These results indicate that the subicular neuron subtypes (burst firing and interneurons) show different patterns of epileptiform discharges with neuronal firing frequency higher in interneurons than in burst firing with 4AP-0Mg perfusion.

As both burst firing and interneurons exhibited different electrophysiological properties before and during epileptogenic discharges (Fig. 1; *SI Appendix* Fig. S2; *SI Appendix* Table S2), the neuronal data from these subicular subtypes were taken for further studies separately to study the modulation of epileptic activity in these two classes of neurons.

### The Intrinsic Sag Response Was Differentially Regulated in Burst Firing Neurons and Interneurons with the HCN Modulation using the Dynamic Clamp

In our study, to examine the functional outcome of alterations in I_h_ in subicular neurons, we artificially manipulated HCN currents using the dynamic clamp in our current-clamp experiments that served as a better alternative to the blockade of I_h_ by pharmacological agents. To ensure the successful incorporation of the subicular I_h_ model in the dynamic clamp system that can mimic the upregulation and downregulation of HCN expressions, we added (+5nS G_h_) and subtracted (–5nS G_h_) HCN currents through the dynamic clamp and studied the sag profile in response to the hyperpolarizing step pulse of –1 nA in a burst firing neuron. The added HCN current resulted in increased sag with respect to the control voltage sag response (Fig. 2*Ba, b*), whereas the subtracted HCN showed a decreased sag compared to the control response (Fig. 2*Ba, c*).

**Figure 2.**
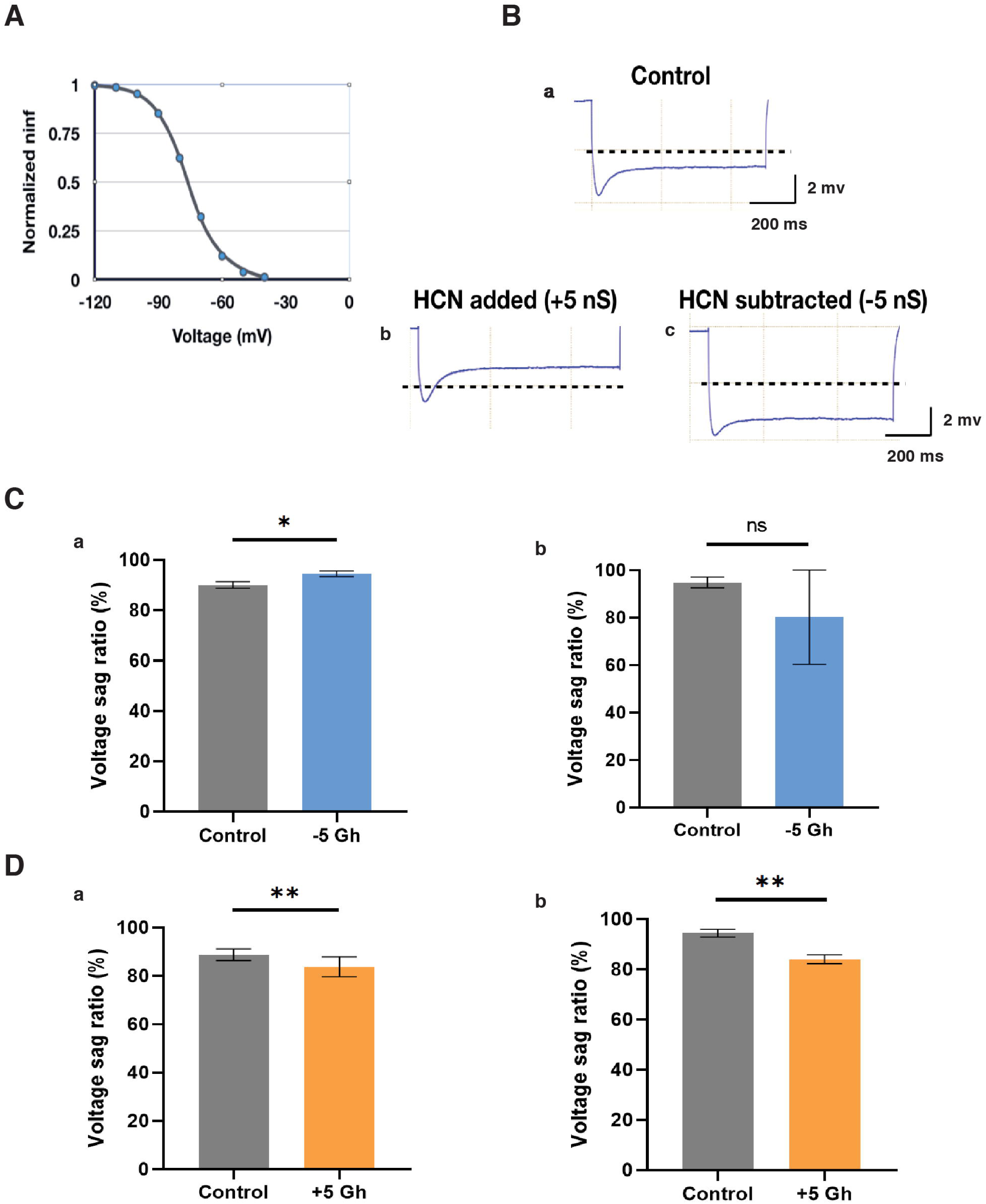
Dynamic clamp-mediated HCN conductance modulation resulted in differential sag responses between burst firing neurons and interneurons. (A) The conductance was modeled from HCN conductance parameters (Eqn.2, Methods) used by van Welie *et al*., 2006 for rat subicular neurons. The channel is activated at hyperpolarized range. (B) (a) The voltage response to –1 nA hyperpolarizing pulse with characteristic sag; (b) The addition of I_h_ (+5nS) increases the sag; (c) The subtraction of I_h_ (–5nS) decreases the sag. The dashed line is a reference voltage at –68 mV. (C) Changes in sag ratio (%) under control conditions and after activating the artificial I_h_ with G_h_ = –5nS through the dynamic clamp in (a) burst firing neurons (control 90.06 ± 1.32 vs –5nS G_h_ 94.48 ± 1.15, n=5, * p <0.5, paired t-test) and (b) in interneurons (control 94.78 ± 2.28 vs –5nS G_h_ 80.2 ± 19.8, n=5, ns p >0.5, paired t-test). (D) Changes in sag ratio (%) under control conditions and after adding the artificial I_h_ with G_h_ = +5nS through dynamic clamp (a) in burst firing neurons (control 88.71 ± 1.07 vs +5nS G_h_ 83.72 ± 1.84, n=8, ** p <0.05, paired t-test) and (b) in interneurons (control 94.41 ± 1.54 vs +5nS G_h_ 83.99 ± 1.82, n=8, ** p <0.05, paired t-test). Data represent mean ± SEM in all plots.

The sag ratio, calculated as the difference between the steady state and the peak voltage in response to a hyperpolarizing step pulse, is a characteristic of HCN channel (I_h_) activation (26–28). We studied the percentage sag ratio to the hyperpolarizing current pulse after activating artificial I_h_ with G_h_ = –5nS and G_h_ = +5nS for the subtraction and addition of artificial I_h,_ respectively (Fig. 2*C-D*). This was done in both burst firing neurons and interneurons to observe if there are differences in HCN activity in these two classes of subicular neurons. We observed a significant increase in the percentage sag ratio in control conditions and after subtracting the artificial I_h_ with G_h_ = –5nS through the dynamic clamp in burst firing neurons (control 90.06 ± 1.32 vs –5nS G_h_ 94.48 ± 1.15, n=5, * p <0.5, paired t-test) (Fig. 2*Ca*). However, in interneurons, there were no changes in percentage sag ratio after the activation of artificial –I_h_ of similar conductance (control 94.78 ± 2.28 vs –5nS G_h_ 80.2 ± 19.8, n=5, ns p >0.5, paired t-test) (Fig. 2*Cb*).

Similarly, we studied the percentage sag ratio profile by adding artificial I_h_ G_h_ = +5nS in burst firing and interneurons. We observed a significant change in the percentage sag profile after the addition of artificial I_h_ through the dynamic clamp in both burst firing neurons (control 88.71 ± 1.07 vs +5nS G_h_ 83.72 ± 1.84, n=8, ** p <0.05, paired t-test) (Fig. 2*Da*) and interneurons (control 94.41 ± 1.54 vs +5nS G_h_ 83.99 ± 1.82, n=8, ** p <0.05, paired t-test) (Fig. 2*Db*). We observed significant changes in the sag activity in both the classes of neurons upon adding I_h_ by dynamic clamp, whereas the subtraction of I_h_ had a significant effect on the sag function in burster cells but showed no changes in interneurons.

### 4AP 0Mg-Induced Epileptogenic Activity can be Modulated by HCN Conductance Manipulation in Burst Firing Neurons but Not in Interneurons

Alterations in I_h_ can reflect both hyperexcitability (9, 13) and reduced excitability (16, 29) arising from its physiological conditions. Using the dynamic clamp, we investigated how I_h_ modulates epileptiform firing in contexts of both increased and decreased neuronal excitability. After characterizing burster cells and interneurons followed by the establishment of epileptiform activity, the control recording was obtained with nACSF solution with the perfusion of 4AP-0Mg for 30s followed by activating artificial I_h_ with the G_h_ = –5nS through the dynamic clamp for 30s and inactivating I_h_ with G_h_ = 0nS for the following 30s in both burst firing cells and interneurons. The firing activity with 4AP-0Mg is considered as control in all the recordings in these experiments.

In the burst firing neurons, the firing rate with the activation of artificial –I_h_ showed a significant increase from that observed in the control 4AP-0Mg environment (control 6.35 ± 1.64 Hz vs – 5nS G_h_ 11.25 ± 1.79 Hz, n=4, ** p <0.01, paired t-test), whereas the inactivation of I_h_ with G_h_= 0 nS brought back the firing activity as that in control environment (–5nS G_h_ 11.25 ± 1.79 and 0nS G_h_ 6.12 ± 1.39 Hz, n=4, ** p <0.01, paired t-test) (Fig. 3*Ca*). We further examined the modulation of epileptiform activity with the addition of artificial I_h_ in these burst firing neurons. The firing rate with the activation of artificial +I_h_ showed a significant decrease from that observed in the control 4AP-0Mg environment (control 8.12 ± 4.04 Hz vs +5nS G_h_ 5.75 ± 3.77 Hz, n=4, ** p <0.01, paired t-test), whereas the inactivation of I_h_ +5nS G_h_ i.e., I_h_ with G_h_= 0 nS brought back the firing activity as that in control conditions (+5nS G_h_ 5.75 ± 3.77 Hz and 0nS G_h_ 7.85 ± 3.78 Hz, n=4, ** p <0.01, paired t-test) (Fig. 3*Cb*).

**Figure 3.**
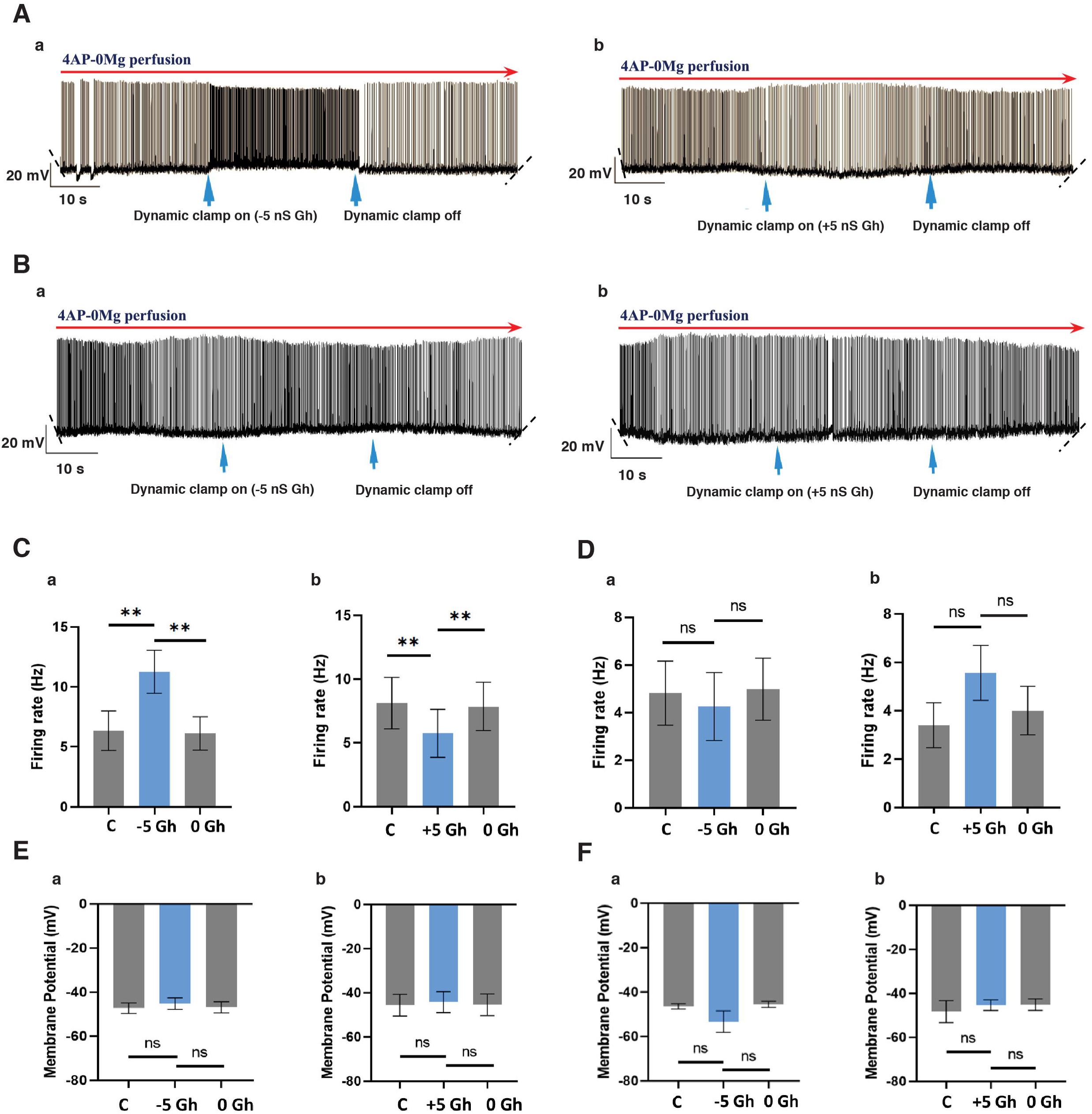
Modulation of HCN using the dynamic clamp regulate 4AP 0Mg-induced epileptogenic firing in burster cells, whereas it showed no effect in interneurons. (A) Voltage trace showing epileptiform activity using 4AP-0Mg (30s) followed by the activation of artificial I_h_ through the dynamic clamp that was turned on (30s) and off (30s) at the arrows (blue) in a burst firing neuron with (a) –5nS G_h_ (b) +5nS G_h._ (B) Voltage trace showing epileptiform activity using 4AP-0Mg (30s) followed by the activation of artificial I_h_ through dynamic clamp that was turned on (30s) and off (30s) at the arrows (blue) in an interneuron with (a) –5nS G_h_ (b) +5nS G_h._ (C) Histogram showing the mean firing rate in burst firing neurons during 4AP-0Mg induced epileptiform activity for 30s followed by the activation of (a) – 5nS G_h_ for 30s in (control 6.35 ± 1.64 Hz vs –5nS G_h_ 11.25 ± 1.79 Hz, n=4, ** p <0.01, paired t-test), followed by the inactivation of –5nS G_h_ for 30s (–5nS G_h_ 11.25 ± 1.79 Hz and 0nS G_h_ 6.12 ± 1.39 Hz, n=4, ** p <0.01, paired t-test) and (b) +5nS G_h_ for 30s (control 8.12 ± 4.04 Hz vs +5nS G_h_ 5.75 ± 3.77 Hz, n=4, ** p <0.01, paired t-test), followed by the inactivation of +5nS G_h_ for 30s (+5nS G_h_ 5.75 ± 3.77 Hz and 0nS G_h_ 7.85 ± 3.78 Hz, n=4, ** p <0.01, paired t-test). (D) Profile of the mean firing rate in interneurons during 4AP-0Mg induced epileptiform activity for 30s followed by the activation of (a) –5nS G_h_ for 30s (control 4.81 ± 1.34 Hz vs –5nS G_h_ 4.25 ± 1.43 Hz, n=6, ns p = 0.33, paired t-test) followed by inactivation of –5nS G_h_ for 30s (–5nS G_h_ 4.25 ± 1.43 Hz vs 0nS G_h_ 4.98 ± 1.3 Hz, n=6, ns p = 0.18, paired t-test) (b) +5nS G_h_ for 30s (control 3.4 ± 0.92 Hz vs +5nS G_h_ 3.56 ± 1.14 Hz, n=5, ns p = 0.16, paired t-test), followed by the inactivation of +5nS G_h_ for 30s (+5nS G_h_ 3.56 ± 1.14 Hz vs 0nS G_h_ 4 ± 1.0 Hz, n=5, ns p=0.07, paired t-test). (E) Summary plot of membrane potentials in burst firing neurons during 4AP-0Mg induced epileptiform activity for 30s followed by the activation of (a) –5nS G_h_ for 30s in (control –47.29 ± 2.38 mV vs –5nS G_h_ –45.24 ± 2.6 mV, n=4, ns p =0.29, paired t-test), followed by the inactivation of –5nS G_h_ for 30s (–5nS G_h_ –45.24 ± 2.6 mV vs 0nS G_h_ –46.93 ± 2.48 mV, n=4, ns p =0.17, paired t-test) (b) +5nS G_h_ for 30s (control –43.67 ± 4.92 mV vs +5nS G_h_ –44.26 ± 4.78 mV, n=4, ns p =0.17, paired t-test), followed by the inactivation of +5nS G_h_ for 30s (+5nS G_h_ – 44.26 ± 4.78 mV vs 0nS G_h_ –45.47 ± 4.9 mV, n=4, ns p =0.3, paired t-test). (F) Changes in membrane potentials in interneurons during 4AP-0Mg induced epileptiform activity for 30s followed by the activation of (a) –5nS G_h_ for 30s (control –46.5 ± 1.23 mV vs –5nS G_h_ –53.36 ± 4.88 mV, n=6, ns p =0.23, paired t-test), followed by the inactivation of –5nS G_h_ for 30s (–5nS G_h_ –53.36 ± 4.88 mV vs 0nS G_h_ –45.55 ± 1.37 mV, n=6, ns p =0.17, paired t-test) (b) +5nS G_h_ for 30s (–48.32 ± 2.21 mV vs +5nS G_h_ –45.4 ± 1.07 mV, n=5, ns p = 0.18, paired t-test) followed by inactivation of +5nS G_h_ for 30s (+5nS G_h_ –45.4 ± 1.07 mV vs 0nS G_h_ max –45.16 ± 1.17 mV, n=4, ns p =0.79, paired t-test). Data represent mean ± SEM in all plots.

However, in interneurons, the addition and subtraction of artificial I_h_ through the dynamic clamp showed no detectable changes in their firing frequency. We observed that the firing rate after activating the artificial I_h_ with G_h_ = –5nS through the dynamic clamp (control 4.81 ± 1.34 Hz vs –5nS G_h_ 4.25 ± 1.43 Hz, n=6, ns p = 0.33, paired t-test) was not significant after which the firing activity with the inactivation of I_h_ = –5 nS, i.e., I_h_ with G_h_= 0 nS also did not show any significant changes from that observed under activated I_h_ (–5nS G_h_ 4.25 ± 1.43 Hz vs 0nS G_h_ 4.98 ± 1.3 Hz, n=6, ns p = 0.18, paired t-test) (Fig. 3*Da*).

Similarly, we observed no significant changes in the firing rate after activating and inactivating the artificial I_h_ with +5nS G_h_ through the dynamic clamp (activating I_h_: control 3.4 ± 0.92 Hz vs +5nS G_h_ 3.56 ± 1.14 Hz, n=5, ns p = 0.16, paired t-test; and inactivating I_h_: +5nS G_h_ i.e., I_h_ with G_h_= 0 nS 3.56 ± 1.14 Hz vs 0nS G_h_ 4 ± 1.0 Hz, n=5, ns p=0.07, paired t-test) (Fig. 3*Db*).

Changes in neuronal membrane potential during HCN modulation were studied to understand if I_h_ brought any hyperpolarizing or depolarizing effects during epileptiform activity in burst firing neurons and interneurons. We studied the membrane potentials during control conditions (4AP-0Mg), and during the activation of artificial +I_h_ and –I_h_ through the dynamic clamp followed by their inactivation. In burster neurons, we observed that there were no significant changes in the membrane potentials when negative G_h_ conductance (–5nS) was inserted (control –47.29 ± 2.38 mV vs –5nS G_h_ –45.24 ± 2.6 mV, n=4, ns p =0.29, paired t-test) and then inactivated (–5nS G_h_ –45.24 ± 2.6 mV and 0nS G_h_ –46.93 ± 2.48 mV, n=4, ns p =0.3, paired t-test) during the epileptiform activity (Fig. 3*Ea*). There were no significant changes also with the addition of I_h_ conductance (+5nS) (control –43.67 ± 4.92 mV vs +5nS G_h_ –44.26 ± 4.78 mV, n=4, ns p =0.17, paired t-test) and when the dynamic clamp was turned off (+5nS G_h_ – 44.26 ± 4.78 mV vs 0nS G_h_ –45.47 ± 4.98 mV, n=4, ns p =0.17, paired t-test) during 4AP 0Mg-induced epileptiform activity (Fig. 3*Eb*). In interneurons, we observed that there were no significant changes in the membrane potentials when negative G_h_ conductance (–5nS) was inserted and removed (control –46.5 ± 1.23 mV vs –5nS G_h_ –53.36 ± 4.88 mV, n=6, ns p =0.23, paired t-test; –5nS G_h_ –53.36 ± 4.88 mV vs 0nS G_h_ –45.55 ± 1.37 mV, n=6, ns p =0.17, paired t-test) (Fig. 3*Fa*). There were no significant changes also with the addition of G_h_ conductance (+5nS) control –48.32 ± 2.21 mV vs +5nS G_h_ –45.4 ± 1.07 mV, n=5, ns p = 0.18, paired t-test; +5nS G_h_ –45.4 ± 1.07 mV vs 0nS G_h_ –45.16 ± 1.17 mV, n=5, ns p =0.79, paired t-test (Fig. 3*Fb*).

I_h_ contributes differently to modulating epileptiform activity in two different classes of neurons in the subiculum. We observed that the burst firing neurons primarily contribute to the characteristics of epileptogenic firing due to HCN in the subiculum, whereas interneurons did not respond to changes in HCN conductance during the epileptiform activity.

### Rapid Progressive Loss of Homeostatic Modulation of HCN Activity was Observed during 4AP 0Mg-Induced Epileptiform Activity in Subicular Burst Firing Neurons

To evaluate how the subicular burst firing neurons contribute to the epileptic firing characteristics due to HCN in the subiculum, we studied the homeostatic modulation of HCN activity in the subicular neuron during 4AP 0Mg-induced epileptiform activity. Fig. 4*(A-B)* is an illustration of changes in the membrane potential to hyperpolarizing step current injections from –350 pA to –10 pA with 20 pA incremental steps for 1s to study the changes in sag profile during epileptiform activity induced by 4AP-0Mg.

**Figure 4.**
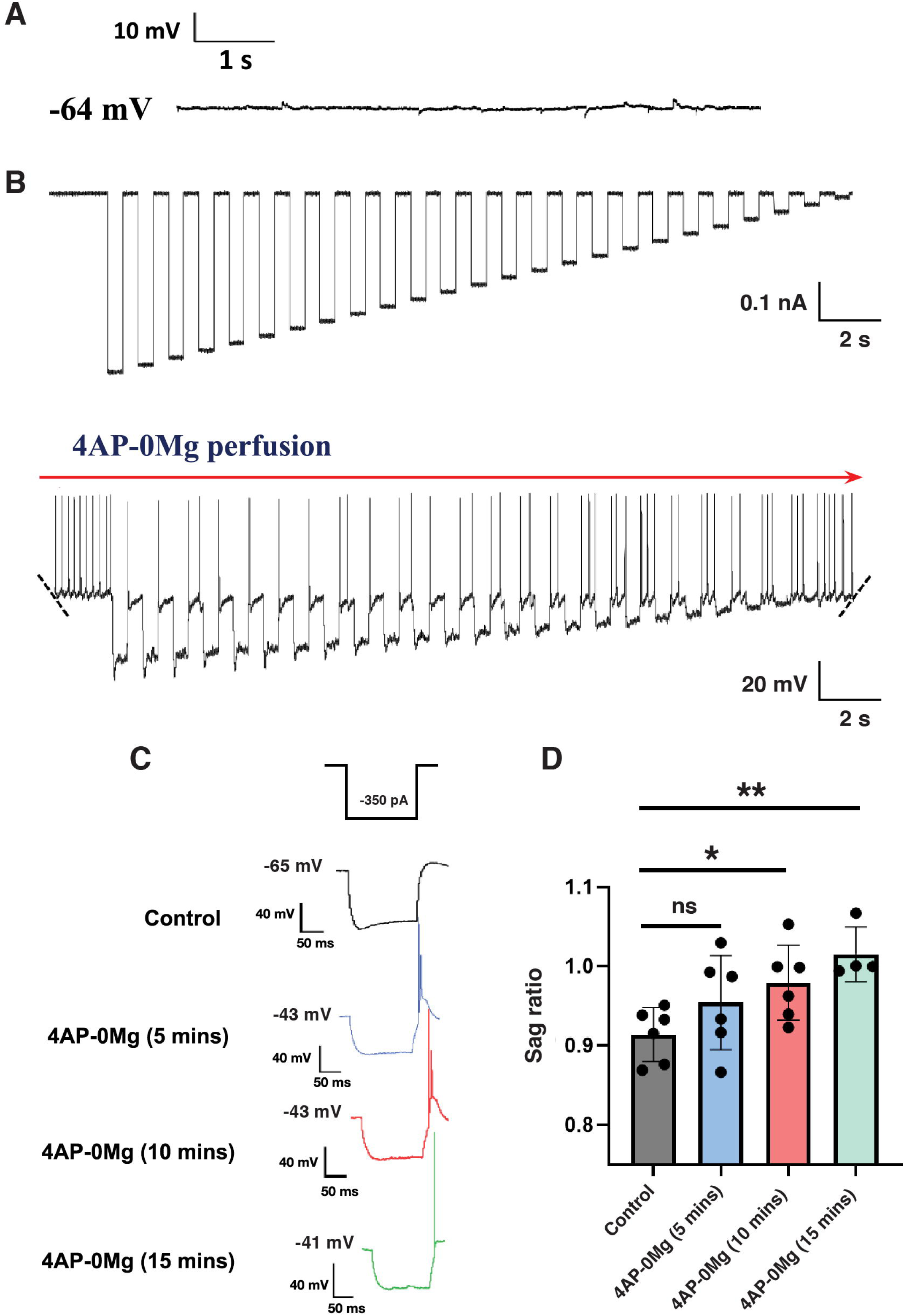
Quantitative analyses of the sag ratio at different timepoints during the epileptiform activity in burst firing neurons show rapid modulation of HCN activity within minutes. (A) Recording trace of the basal activity at RMP = –64 mV in the burst firing neuron. (B) Injection of hyperpolarizing step currents from –350 pA to –10 pA for 1s with an incremental step of 20 pA to study sag responses during the epileptiform activity induced by 100 μM 4AP-0Mg (top) and their subsequent voltage response (bottom). The membrane potential of the cell was – 46.7 mV. (C) An example of voltage waveforms in response to the hyperpolarizing step current –350 pA for 1s in control conditions and at different timepoints (5 mins; 10 mins; 15 mins) after induction of epileptiform activity with 4AP-0Mg perfusion. (D) Histogram of changes in the sag ratio in control conditions and at different timepoints (5 mins; 10 mins; 15 mins), following the induction of epileptiform activity by 4AP-0Mg perfusion (control vs 4AP-0Mg (5 mins), n=6, ns p=0.17; control vs 4AP-0Mg (10 mins), n=6, *p<0.05; control vs 4AP-0Mg (n=4) (15 mins), **p<0.01; unpaired t-test). Data are represented as mean ± SEM.

The changes in the HCN channel activity with time were studied from the sag response to step current injections of –350 pA for 1s during different timepoints *viz*. 5, 10, and 15 mins after 4AP-0Mg perfusion. The sag ratio was analyzed in the control conditions and at different timepoints during the epileptiform activity that displayed the progression of HCN activity during the epileptiform activity over a short time frame (Fig. 4*C*).

It was observed that the profile of the sag ratio in control conditions and at the onset of epileptiform activity with 4AP-0Mg did not show any significant change (control 0.91 ± 0.01 and 4AP (5 mins) 0.95 ± 0.02, n=6, ns p=0.17, unpaired t-test). However, following the induction of epileptiform activity and its temporal progression, there was a significant change in the sag ratio from control and 4AP-0Mg (10 mins) (control 0.91 ± 0.01 and 4AP-0Mg (10 mins) 0.97 ± 0.01, n=6, * p<0.05, unpaired t-test). These changes in the sag ratio further increased after 5 mins that were significant (control 0.91 ± 0.01 and 4AP-0Mg (15 mins) 1.0 ± 0.01, n=4, ** p<0.01, unpaired t-test) (Fig. 4*D*). These results indicate a rapid progressive loss of HCN-associated compensation during the 4AP-0Mg epileptiform activity.

### Epileptiform Activity-Induced Alterations in HCN Activity Are Irreversible post 4AP-0Mg Washout, Leading to Hyperexcitability in Burst Firing Neurons

We studied if the changes induced by the epileptiform activity in these burst firing neurons are reversible or irreversible after 4AP-0Mg washout. We investigated these changes by analyzing the sag ratio and the chirp stimulus responses in two environmental conditions, i.e., control (before 4AP-0Mg perfusion) and 5 mins after the 4AP-0Mg washout. Changes in the sag ratio were studied by the voltage responses elicited from the hyperpolarizing current of –350 pA for 1s, and the changes in resonance amplitude responses were studied by injecting a chirp stimulus. Both the stimuli were given at the RMPs of the neurons under control and 4AP-0Mg washout conditions. Fig. 5 shows the results obtained in the study conducted to observe these intrinsic changes in the neurons. We observed that the sag ratio in control and 4AP-0Mg washout conditions showed a significant change (control 0.91 ± 0.01 and 4AP-0Mg washout 0.96 ± 0.01, n=6, * p<0.05, paired t-test) (Fig. 5*A*, *C*). The resonance frequency responses in control at RMP = –65 mV and 4AP-0Mg washout at RMP = –53 mV also showed a significant change (control 4.01 ± 0.01 Hz and 4AP-0Mg washout 1.64 ± 0.06 Hz, n=6, *p<0.05, paired t-test) (Fig. 5*B*, *D*). These changes in the sag ratio and resonance frequency profile showed irreversible epileptiform-induced alterations in the HCN-directed intrinsic properties of the neurons post 4AP-0Mg washout.

**Figure 5.**
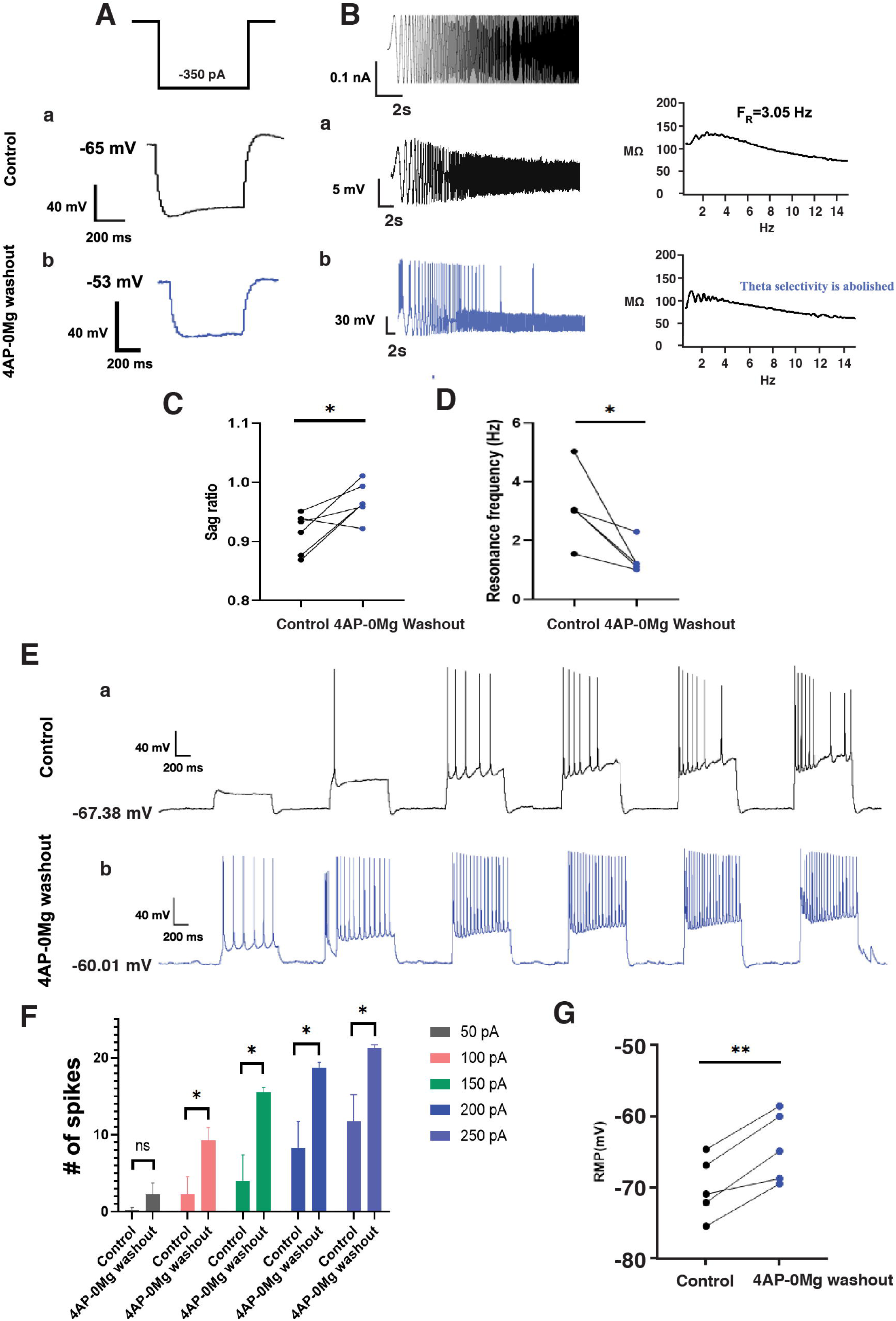
Irreversible changes in the sag ratio and resonance response observed after 4AP-0Mg washout in burst firing neurons resulted in subsequent neuronal hyperexcitability. (A) Voltage response to the hyperpolarizing step current –350 pA for 1s (a) in control conditions and (b) after the 4AP-0Mg washout. (B) A chirp stimulus evoked different voltage responses (a) in control conditions (black; RMP=-65 mV) and (b) after the 4AP-0Mg washout (blue; RMP=-53 mV). The respective impedance amplitude profiles of neurons in two experimental conditions (a, b) are represented to the right of their voltage-frequency response. (C) Profile of the sag ratio from control (black) and after the washout of 4AP-0Mg (blue) from 6 neurons (n=6; *p<0.05, paired t-test). (D) Changes in resonance frequency response from 6 neurons in control (black; RMP=-65 mV) and 4AP-0Mg washout (blue; RMP=-53 mV) conditions (n=5; *p<0.05, paired t-test). (E) Voltage trace of the spiking activity of a burst firing neuron in response to graded depolarizing step currents from 50 pA to 300 pA with the incremental steps of 50 pA for 1s (a) in control and (b) 4AP-0Mg washout conditions. (F) Average data of change in number of spikes in control and 4AP-0Mg washout conditions as a function of depolarizing currents for 1s with amplitudes 50 pA (control 0.25 ± 0.25 and 4AP-0Mg washout 2.25 ± 1.43, n=4, ns p= 0.2192; 100 pA (control 2.25 ± 2.25 and 4AP-0Mg washout 9.25 ± 1.65, n=4, *p <0.05; 150 pA (control 4 ± 3.34 and 4AP-0Mg washout 15.5 ± 0.64, n=4, *p <0.05); 200 pA (control 8.25 ± 3.44 and 4AP-0Mg washout 18.75 ± 0.62, n=4, *p =<0.05); 250 pA (control 11.75 ± 3.47 and 4AP-0Mg washout 21.25 ± 0.47, n=4,*p<0.05); unpaired t-test. (G) RMP changes of the burst firing neurons before (black) and after (blue) the epileptiform activity (n=5, **, p<0.01, paired t-test). Data are represented as mean ± SEM.

We further investigated how these epileptiform activity-induced irreversible changes in HCN function affect the overall excitability of the neurons. We studied the spiking activity of burst firing neurons before and after the 4AP 0Mg-induced epileptiform activity by applying graded depolarizing step currents from 50 pA to 250 pA with incremental steps of 50 pA for 1s to test the epileptiform-induced modifications in the excitability of the neurons. The average data of change in the number of spikes as a function of depolarizing currents for 1s is given in Fig. 5*F*. We saw a tendency for higher spiking of neurons after the termination of the epileptiform activity by 4AP-0Mg washout. The number of spikes evoked in response to 50 pA did not change significantly after the 4AP-0Mg washout (control 0.25 ± 0.25 and 4AP-0Mg washout 2.25 ± 1.43, n=4, ns p = 0.21, unpaired t-test). However, there were significant changes in the number of spikes elicited in response to 100 – 250 pA (100 pA (control 2.25 ± 2.25 and 4AP-0Mg washout 9.25 ± 1.65, n=4, *p <0.05); 150 pA (control 4 ± 3.34 and 4AP-0Mg washout 15.5 ± 0.64, n=4, *p <0.05); 200 pA (control 8.25 ± 3.44 and 4AP-0Mg washout 18.75 ± 0.62, n=4,* p <0.05); 250 pA (control 11.75 ± 3.47 and 4AP-0Mg washout 21.25 ± 0.47, n=4,*p <0.05), unpaired t-test) (Fig. 5*F*).

We also studied the resting membrane potentials (RMPs) of these neurons before and after the epileptiform activity. The RMP changes were monitored under control environment and after 4AP-0Mg washout and showed a significant change in the RMP at these two conditions (control –69.95 ±1.92 mV and 4AP-0Mg washout –64.25 ± 2.23 mV, n=4, **p <0.01, paired t-test) (Fig. 5*G*).

The epileptiform-induced changes in the sag and resonance frequencies in the neurons that we observed earlier indicate an acquired alteration in the HCN channel function. These changes, in turn, produce an increase in the intrinsic excitability of individual neurons (Fig. 5*E-G*) that can lead to network hyperexcitability that can, in turn, lead to recurrent seizures.

## DISCUSSION

TLE has been associated with changes in HCN channel expressions (16, 30); however, there have been conflicting observations from animal models showing that I_h_ can be upregulated or downregulated in epileptic conditions depending on the physiological context (9, 22). In the current work, the contribution of I_h_ in subicular neuronal subtypes to shape epileptiform firing was studied by artificially manipulating I_h_ through a dynamic clamp that could mimic the upregulation and downregulation of HCN channels observed in epileptic conditions. We observed that the 4AP 0Mg-induced epileptogenic activity can be modulated by HCN conductance manipulation in burst firing neurons but does not cause any change in interneurons (Fig. 3). Our experiments imply that I_h_ contributes differently to modulating epileptiform activity in two distinct classes of neurons in the subiculum and the differential response of I_h_ in these neuronal subtypes could influence the pattern of the epileptogenic processes in the subiculum. Our study suggests that the burst firing neurons determine the characteristics of the epileptiform firing due to HCN in the region of subiculum. Thus, I_h_ endows the burster cells with properties that favour epileptic firing, whereas it does not have a significant effect on interneurons. Our results show that burst firing neurons have a direct link between HCN modulation and epileptiform activity. Reports have demonstrated that the bursters may initiate the focal epileptiform activity in the subiculum (31), and our study proposes that I_h_ is one of the significant intrinsic factors in burst firing neurons that may aid in initiating epileptogenesis in the subiculum.

On the contrary, interneurons not being affected by HCN modulation, unlike the burst firing neurons, is an interesting phenomenon. This can be because of different degrees of sag (32, 33) in these two types of neurons, as also observed in our study, that reflect their different HCN current densities that lead to varied functional impact of I_h_ in them. In non-pyramidal inhibitory neurons, HCN currents are primarily localized in the soma and, hence, its effect on the neuronal excitability is carried out via its depolarized membrane potential, leading to the opposite excitatory effect in interneurons than what is observed in principal neurons (34). As HCN interplay with other subthreshold conductances like persistent sodium channels (35), T-type calcium channels (36–37) and M-type calcium channels (38–40), further investigations can examine if some ion channel or signaling cascade would be required to bring about a change in epileptiform characteristics due to I_h_ in interneurons.

In the epileptic brain, various cellular alterations that occur during epileptiform activities result in increased excitability that is not compensated by homeostatic plasticity. Studies have indicated that a more rapid mechanism of plasticity at the cellular level exists (41–42). Such mechanism of plasticity includes the rapid modulation of I_h_, which contributes to neuronal plasticity occurring within minutes (32, 43), and this rapid upregulation attributed to HCN1 channel trafficking and surface expression (44).

However, the involvement of homeostatic compensation at the cellular level in epileptogenic processes remains largely unexplored. Furthermore, the alterations in homeostatic responses of voltage-dependent conductances during epileptogenic activity, as well as the timeframe over which these alterations occur, have not been adequately investigated. We examined HCN-directed homeostatic response that reflects I_h_-associated cellular plasticity during 4AP 0Mg-induced epileptiform activity. The homeostatic response of HCN activity in which the sag function was preserved for the initial few minutes was later observed to undergo a progressive loss in the next few minutes of the epileptiform activity. Our results indicate that HCN activity in rat subiculum neurons during 4AP 0Mg-induced epileptiform activity is subject to modulation in a timeframe of a few minutes (Fig. 4*C, D*). Our study is an early report that shows that the HCN-associated homeostatic response in burst firing neurons, following the induction of epileptiform activity, undergoes a progressive decline over a very short time course in the region of subiculum. The gradual loss in the homeostatic response of HCN activity in epileptogenesis is indicative of the failure of cellular homeostasis to counteract these alterations that can lead the neuron’s trajectory towards seizure space. While studies have demonstrated rapid increase in HCN function over short time scale (32), investigations observing decreased HCN activity within the same time span, reflecting potential deficits in homeostatic plasticity in regulating early neuronal and network hyperexcitability in epileptogenesis, are not yet conducted. Our result is also the first report that shows decreased activity of HCN in a neuron within a time span of a few minutes.

In our study, we have investigated whether the changes observed in HCN activity during 4AP-0Mg induced epileptiform activity were reversible or irreversible post 4AP-0Mg washout. The persistent decreased HCN function after wash-out of 4AP-0Mg suggest that these alterations are irreversible (Fig. 5*C, D*). Therefore, our data demonstrate that these epileptiform activities can modify intrinsic HCN function, indicating acquired proepileptic channel characteristics in the subicular bursting cells. These modulations that are irreversible after 4AP-0Mg washout can lead to increased excitability in the subiculum burst firing neurons, which we have studied further (Fig. 5*E-G*). Studies suggest seizure-induced changes in the immature brain can result in irreversible alterations in neuronal connectivity (45, 46). Younger animals show faster seizure onset, leading to rapid kindling and widespread seizures (47). Our study is an attempt to understand the seizure mechanisms in immature neurons and contribute insights for developing treatments in the younger and juvenile populations.

The phenomenon involving HCN function and expression alterations progresses to a state of increased excitability of neurons and consequent epileptogenesis (16, 48). HCN1, abundantly found in the subiculum (49), has been implicated in hyperexcitability within cortical (50), and hippocampal regions (51) as observed in these HCN1 knockout studies and has shown prolonged excitatory responses to synaptic stimulation (50). We further studied the spiking activity of burst firing neurons before and after the 4AP 0Mg-induced epileptiform activity to examine the effect of epileptiform-induced modifications on the intrinsic excitability of the neurons. We observed neurons exhibiting higher spiking activity after the termination of epileptiform activity by 4AP-0Mg washout. This acquired hyperexcitability observed in our study in the region of subiculum and several other studies depicting enhanced excitability in the hippocampal regions across various animal models in the epileptogenic processes are in contrast with the HCN activity observed in the thalamic region during absence seizures (AS). It has previously been shown that blocking I_h_ in the ventrobasal thalamus (VB) led to the suppression of burst firing and eliminated spontaneous absence seizures, whereas genetic knockdown of HCN decreased absence seizures (52, 53). Therefore, it is suggested that any potential therapy for TLE must take into account the opposite role of hippocampal and thalamic HCN currents.

The spatial arrangement of HCN channels within dendritic compartments of subicular pyramidal neurons is 60-fold greater than that of the somatic HCN channels (49). Examining whether dendritic HCN channels undergo similar modulation as somatic HCN channels, potentially eliciting pro-epileptic effects during epileptogenesis or exhibiting contrasting responses leading to compensatory homeostatic mechanisms, would be of considerable interest.

## ACKNOWLEDGMENTS

We thank DST-INSPIRE, Govt. of India, for funding Monica Alfred during her PhD. The research was partially funded by the DBT-IISc Partnership Program for Advanced Research in Biological Sciences and Bioengineering (DBT, India).

## AUTHOR CONTRIBUTIONS

M.A. and S.K.S. designed the research; M.A. performed the experiments and analyzed data; and M.A. and S.K.S. interpreted the data and wrote the paper.

## DECLARATION OF INTERESTS

The authors declare no competing interests.

## Notes

### Competing Interest Statement

The authors have declared no competing interest.

