## Supplementary Information for "Role of somatic HCN in epileptiform activity in subicular neurons"

---

### The Recording Configuration in the Dynamic Clamp System

The connections for the channels described below were made in the following configuration between the amplifier (Dagan BVC 700A) and the data acquisition board (NI-PCI 6221):

1. Dagan BVC 700A Vm output => DAQ Analog Input 1

2. Dagan BVC 700A I output => DAQ Analog Input 0

3. Dagan BVC 700A External Command Input => DAQ Analog Output 0

The calibration is done using the following command:
"$ sudo comedisoftcalibrate"

The command "sudo comedisoftcalibrate" was used to calibrate the DAQ card after connecting the appropriate channels. The total gain calculated by the amplifier (Dagan BVC 700A) was computed. Subsequently, the scaling factor calculated was fed into the system control panel (Control → System Control).
The gain of the amplifier was set at 10 and the scaling factor for Input 1 in System Control was 1/10/V/V (invert the gain) or 10 mV/V. The amplifier gain for Input 0 (I output) and Output 0 (Ext. Command) were 1000 mV/nA and 0.2 nA/V, respectively. The amplifier gain for these channels was inverted and the estimated scaling factors were given into the respective channels: 1 nA/V for Input 0 and 5GV/A for Output 0. These values were applied to the real-time setup. The real-time sampling rate was set to 10 kHz in the system control. (Ref. Dagan BVC 700A operating manual for details on gain factors).

### Dynamic Clamp Experimentation

The command to initiate the RTXI program in the terminal is by giving the command "sudo rtxi". Access to the digital oscilloscope in RTXI is through: System → Oscilloscope from the RTXI menu bar. Easily customizable modules can be obtained from the RTXI website. Every module can be compiled using the GCC compiler and installed using the command “sudo make install”. Each module will consist of a single class header file (*.h), class implementation file (*.cpp), and a Makefile. All modules are compiled as “*.so” (Linux shared object libraries), that are in link with the core system.

A module can be loaded by selecting the following:

Control → Plugin loader →’plugin name’

The “Connector” is then opened to connect the input-output blocks to an appropriate signal. The module “Data Recorder” in RTXI is used to acquire the data, which is then stored and saved. This can be done using the options: File → Save → “file name”.set. This can be later accessed by following through File → Load →"file name".set.

### Custom Modules and Implementations

Different modules were downloaded from the RTXI website or were written as per the requirement for the dynamic clamp experiments. For the current injection, the current equation as a function of time was specified and injected. For currents specifying ion channels, the driving force given by ($V - E_{rev}$) is calculated and the current is injected into the neuron (1, 2, 3, 4) according to the following equation:

$I = g(V) * (V - E_{rev})$

where, V = membrane potential of the cell

g (V) = ionic conductance that depends on the membrane voltage

$E_{rev}$ = reversal potential of the ionic current

The voltage dependence for the above ion channel equations has to be modeled using either the Boltzmann fit of the steady-state activation curve of the voltage gated ion channel or specifying the rate constants of the HH (Hodgkin-Huxely) parameters (5, 1, 6).

### Checking the Modules in Dynamic Clamp with the Model Cell

The modules customized or written were first tested with a model cell that consisted of 10 MΩ (electrode resistance) in series with a parallel circuit that consisted of 500 MΩ and 6.2 pF, related to the membrane resistance and capacitance of a typical biological cell membrane.

**INa_HH2 (sodium current) and IK_HH2 (potassium current) Modules**

Fast $Na^+$ and K+ currents are responsible for action potentials. Membrane voltage dependent $Na^+$ and K+ currents were modeled using HH equations. These modules were obtained from HH2.mod file (7). They were then adapted in the RTXI format. These conductances were added in the model cell, and their response following their incorporation was tested for the evoked action potential (Fig. S1).

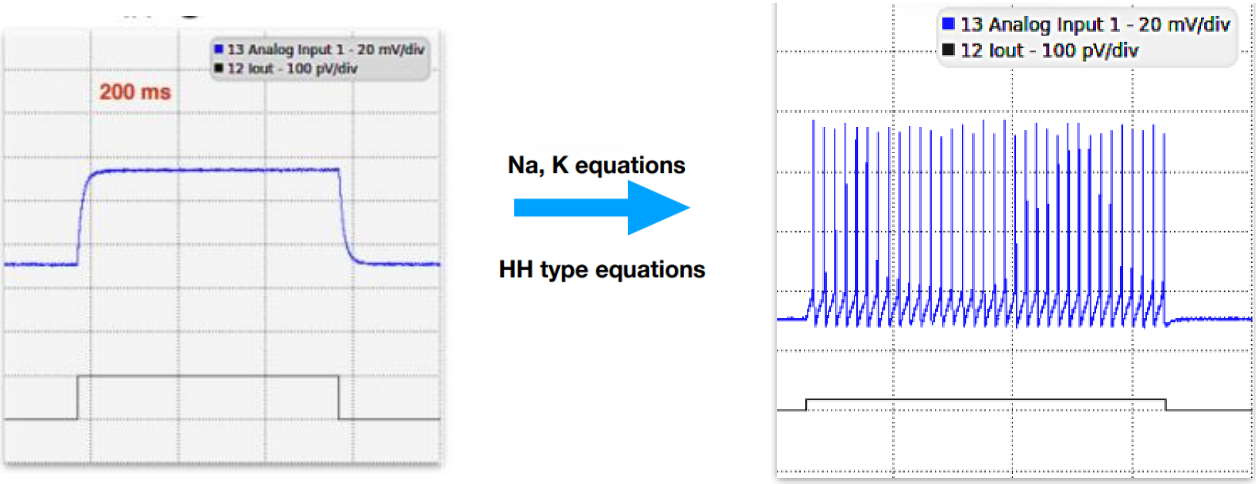


Figure S1. The RC circuit (left), when injected with Na and K currents (HH equations) through the dynamic clamp, produced action potentials (right).

**Data Acquisition and Analyses for Dynamic Clamp**

The dynamic clamp data is acquired in HDF5 file format in RTXI (8). HDF5 is the Hierarchical Data Format Version 5 (HDF5). IGOR pro does not separately identify multiple channel data from the HDF5 data file. In order to distinguish data from multiple channels in HDF5 files, custom codes were used to acquire different channels. Further analyses were carried out using software including Igor Pro and MATLAB. IGOR pro custom code was used for data extraction from HDF5 files with 2 simultaneously recorded channels (current and voltage).

Ionic composition of 4AP-0Mg epileptic induction solution is given below

| **Salt** | **Concentration (mM)** |
|---|---|
| Sodium chloride | 125 |
| Potassium chloride | 2.5 |
| Calcium chloride dihydrate | 2 |
| Sodium bicarbonate | 25 |
| Glucose | 10 |
| 4-Aminopyridine (4-AP) | 0.1 |

Table S1. Ionic composition of 4AP-0Mg epileptic induction solution.

### Calculating the gain of a neuron

F-I slope was calculated by fitting the initial slope where F-I relationships were relatively linear, which gave the gain of the neuron (9).

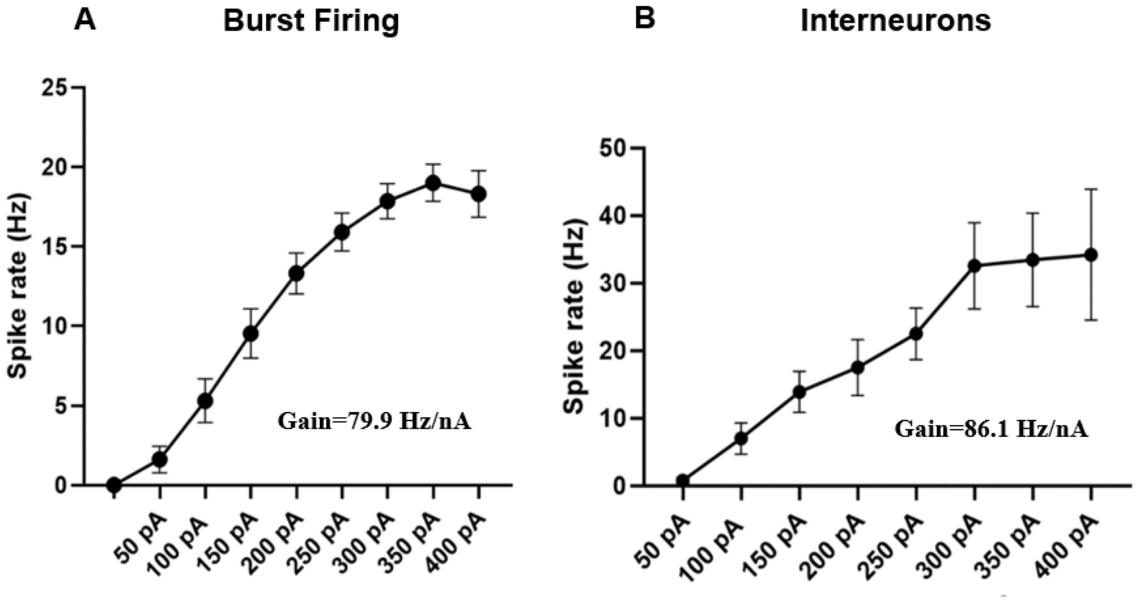


Figure S2. The gain of interneurons is higher than that of burst firing neurons. (A) F-I graph of the burst firing neurons (n=12) showed the gain to be 79.9 Hz/nA (n=12). (B) F-I graph of the interneurons (n=7); the gain was found to be 86.1 Hz/nA (n=7). Data are represented as mean ± SEM.

| **Electrophysiological Properties** | **Burst Firing** | **Interneurons** |
|---|---|---|
| RMP | -66.87 ± 1.19 mV | -70.24 ± 3.16 mV |
| Spike Threshold | -35.91 ± 2.49 mV | -38.33 ± 1.91 mV |
| Spike Amplitude | 66.90 ± 2.74 mV | 55.04 ± 3.5 mV |
| Resonance Frequency | 5.20 ± 1.0 Hz | ** 0.79 ± 0.11 Hz |
| Resonance Amplitude | 167.53 ± 32.53 MΩ | * 293.15 ± 21.68 MΩ |
| Gain | 79.9 Hz/nA | 86.1 Hz/nA |

Table S2. Comparison of different electrophysiological properties of Burst firing and Interneurons.
